# Everything Under Control? Comparing Knepp Estate rewilding project with ‘traditional’ nature conservation

**DOI:** 10.1101/2020.10.12.335877

**Authors:** Benedict Edward Dempsey

## Abstract

‘Rewilding’ is an increasingly prominent concept in conservation, but one that has attracted controversy. Debate frequently focuses on ‘control,’ with rewilding presented as reducing human control of nature. Opposition to rewilding often stems from a perceived lack of control – and associated perception of increased risk and uncertainty.

I explore the concept of control in conservation. I identify that control is not a simple, linear concept, but consists of multiple dimensions. Using a lens of control, I compare two ethnographic case studies: the Sussex Wildlife Trust’s Old Lodge nature reserve; and Knepp Estate, one of the most influential rewilding projects in the UK. These sites ostensibly represent ‘high-control’ and ‘low-control’ examples of conservation.

I outline how Old Lodge does not exert precise control in all respects, but rather involves elements of uncertainty and negotiation. I describe how Knepp’s model of rewilding reduces control in some dimensions but increases it in others. I conclude that, while Knepp’s ‘rewilding’ does represent a significant conceptual departure from ‘traditional’ conservation, it should not be characterised simplistically as an approach that reduces control.

Based on this analysis, I argue that reduction of control should not be assumed to underpin the concept of rewilding. Rather, there is interplay between different control dimensions that combine to form different configurations of control. With this understanding, debate about rewilding – and conservation more broadly – can avoid simplistic characterisations of ‘reducing control’ and become instead an active discussion of what configuration of control is desired.

This analysis could be seen negatively by those who argue that Knepp’s version of rewilding does not sufficiently reduce human control of nature. In contrast, Knepp’s approach can be seen positively as opening new conceptual space while retaining human involvement. It supports the argument that versions of rewilding can be legitimate, innovative components of plural conservation strategies.

## 1. Introduction

The nature conservation community is engaged in a series of fundamental debates about its methods and its purpose. In the context of a global ‘biodiversity crisis’ (1,2) and the widespread assertion that the Earth has entered ‘the Anthropocene’ (3,4), conservationists are being asked to respond with different approaches that meet the challenges of the 21^st^ century (5–7). While these debates differ in different parts of the world they are particularly important in the UK, where they are shaping the creation of new conservation policy frameworks as the country leaves the European Union (EU) (8).

### Rewilding and ‘traditional’ conservation

Within this discussion, ‘rewilding’ has become increasingly prominent, particularly in Europe and North America. While it has no single agreed definition (9), it is often presented as a significant break from modern ‘mainstream’ conservation, eschewing human ‘management’ of nature in favour of greater ‘non-human autonomy’ (10). It is frequently divided into ‘active rewilding’, that involves extensive species (re)introductions to establish dynamic processes; and ‘passive rewilding’ that involves the withdrawal of human intervention to ‘let nature go’ (11). Though varied, rewilding approaches tend to share a shift in focus from the ‘composition’ of an ecosystem’s species assemblage, towards ecosystem functions and processes (12), and the reduction of human management in the long term (13).

Rewilding has been proposed by some as the answer to an array of modern challenges, from mental wellbeing to flood management, as well as addressing the biodiversity crisis (14). By others it has been viewed as a dangerous departure from conservation norms by permitting higher levels of risk (15,16), and a threat to rural communities and ways of life by forcing out traditional farming practices (17).

In this context, there remain questions about what rewilding is and what it represents for conservation, particularly in the UK (18).

### The significance of ‘control’

Within these debates, the issue of ‘control’ is a central concern. ‘Traditional’ forms of European conservation have been described as ‘conservation as control’ that deploy ‘management by interference’ to maintain existing, historical landscapes (19).

Alternative forms of conservation have placed greater emphasis on processes and dynamism – for example, ecological restoration that has attempted to (re)establish functioning dynamic systems (20–22). However, some have argued that conservationists will often only proceed with restoration when they are confident they know the general outcomes of the process – thereby keeping within fixed bounds and attempting control through scientific prediction:

> “We have control not by controlling nature’s every move, but, more cost effectively, by thinking nature’s thoughts”
>
> — (Adams 2003, p.240).

Similarly, approaches like ‘wilderness management’ that seek to preserve ‘pristine’ nature – particularly associated with North America but influential in wider discourse (23) – have been critiqued for representing imagined human conceptions of ‘wildness’ that are nevertheless ‘managed’ (24) and for producing nature that is ‘locked up’ within geographical boundaries and allowed to exist only ‘at the whim of legislators and government policy’ (19).

Recently, the discourse of control in conservation has converged with concepts of biosecurity. Terms such as ‘feral’ are used to describe ‘unsanctioned life’ (such as wild boar) that is challenging and shaping new ideas about human-nature relations, and the control relationship between them (25,26).

The assertion that the Earth has entered the Anthropocene has contributed to suggestions that it is ‘post-wild’ (27) or ‘after nature’ (28). Some, in characterising humanity as if it is a unified entity, have argued that humans have ‘no alternative except to shoulder the mantle of planetary stewardship’ (29). This line of thought implies that uncontrolled, wild nature is a modern impossibility.

In contrast, broader discourse concerning the role of ideas of control in modernity more generally, questions whether notions of ‘control’ are deliverable in anything other than the particular case of well-functioning machines. Stirling (30) argues that when such visions of control (for instance in engineering approaches to Sustainability) meet wider real-world contexts, then ‘control’ is not generally possible. Based on direct routine operational experience with machines, his working definition of control is:

> ‘…realising fully and solely a prior set of intended end(s), with no unintended effects’
>
> — (Stirling, 2019:6)_

On this basis he argues that, in complex contexts, attempts at control are more likely to ‘influence’, ‘impact’ or ‘modify’ some part of the intended ends, rather than effecting ‘control’ over them.

If a term like ‘control’ is held to span such a wide range of meanings in conservation (from ‘influence’ to full machine-like determinism) then this discourse contains, on one side, a sense that control is almost impossible to avoid, and on the other that it is almost impossible to achieve.

### Rewilding and control

For conservation, the idea of ‘rewilding’ brings the concept of control into even sharper focus. For many, rewilding is an explicit attempt to relinquish control, its appeal resting on the proposition that it breaks from excessively controlling, modernistic ‘managerial’ ideas of ‘traditional’ conservation.

Rewilding has been described as ‘resisting the urge to control nature and allowing it to find its own way’ (Monbiot, 2013); as ‘passive management of ecological succession with the goal of restoring natural ecosystem processes and reducing human control of landscapes’ (32); as ‘a process in which a formerly cultivated landscape develops without human control’ (33); and as ‘letting ecosystems evolve out of human control’ (34). Rewilding is often therefore seen to represent a change in the control relationship between people and nature – reducing control and unleashing a nature that is more ‘self-willed’ and embracing unpredicted, emergent properties (35,36).

For Anderson et al (37) ‘…rewilding specifies practices explicitly aimed at a “controlled decontrolling” to promote the agency of nonhuman biophysical processes.’ It is this perception of increased ‘non-human autonomy’ (10), and associated increase in perceived risk, that also underpins much opposition to both ‘passive’ rewilding (33) and ‘active’ rewilding (15).

However, mirroring debates about control more broadly, the degree to which rewilding practice actually delivers ‘low control’ is the subject of continuing debate (18). Jorgensen (9) argues that: ‘the path to this uncontrollable nature is to reinvolve humans and control nature’. Debate includes whether rewilding projects aim to re-establish a form of ‘wild’ nature based on a past baseline (similar to more traditional ecological restoration), versus enabling autonomous processes that proceed into the future without any preconceived ‘end point’ (18,38).

Some have challenged the idea that rewilding delivers non-human autonomy, pointing out the potential contradiction in using ‘autonomous’ non-humans to deliver the human objective of restoration of nature (39). Others have argued that this is too simplistic a view of non-human autonomy, and that rewilding can enable autonomy to emerge in different ways depending on the spatial and political context (40,41). Tension also exists in the rewilding movement between ‘anthropocentric’ and ‘biocentric’ visions – i.e. whether rewilding represents a means to achieve a set of desired ends, or an effort to enable natural autonomy for its own sake (18).

Wynne-Jones et al (18) suggest that, in the UK, the trend among rewilders is away from the recreation of past systems and towards a more novel, future-oriented vision – though activities may be informed by perceived past conditions. They argue that rewilding is inspiring a genuine departure from ‘traditional’ conservation by generating new ideas and conceptual spaces.

They also find, however, that there is so far a limit to which rewilding projects are actually able to ‘let go’ in practice, with a degree of ‘management’ persisting despite ambitions to reduce human intervention. This contributes to uncertainty about the long-term direction of rewilding and the extent to which ‘letting go’ or reduced control is achievable.

### What is control?

‘Control’ is therefore important in debates about conservation generally, and rewilding in particular – but ‘control’ in this context is a complex idea. Despite significant analysis of issues of power and control in conservation (25) it remains a difficult concept to pin down. While others have focused in particular on the political ecology and ‘biopolitics’ of power and control in conservation (18,25,26), this paper seeks to add to the understanding of rewilding’s place in conservation by unpicking what ‘control’ can mean in different settings.

As outlined above, Stirling (30) considers ‘control’ to be ‘…realising fully and solely a prior set of intended end(s), with no unintended effects.’ This machine-inspired characterisation of control can be seen in conservation in a variety of ways.

Perhaps the most straightforward expression of control in conservation might be called ‘stabilisation’. This would describe the types of conservation that Adams (19) calls ‘conservation as control’ – actively intervening to keep an ecosystem stable and prevent it from transitioning into a different form. This has similarities with ‘control theory’ – a concept originating in engineering that uses negative feedbacks to maintain system-stability and prevent deviation, and that has been applied by some to conservation (42). This strand of control relates to wider themes in environmental discourse – including the idea of ‘planetary boundaries’ that compels humanity to maintain nine ‘control variables’ within certain parameters to maintain a ‘safe operating space’ for people (43).

A different apparent strand of control in conservation relates to keeping ‘nature’ within particular geographical boundaries, and could be called control as ‘location’. This type of control appears in Adams’ contention that ‘wild’ nature can be ‘locked up’ in designated places. It relates to debate about ‘land-sparing’ (setting aside specific areas for nature) versus ‘land-sharing’ (making human landscapes more hospitable for biodiversity) – the former indicating elements of control as location (44). It is visible in calls that ‘half the Earth’ should be assigned to nature (45,46). This sort of control could also relate to the distribution of particular habitats and species through concepts and language like ‘non-native’, ‘alien’, ‘invasive’ and ‘feral,’ with the implication that organisms have places that they ought and ought not to be (6,47,48). It is here that conservation and biosecurity potentially converge in the control of ‘unsanctioned life’ (26).

Control may also operate through ‘prediction’. Certain conservation practices might allow dynamic ecological processes, but only permit them to take place if the expected results are considered known and acceptable – described by Adams as ‘thinking nature’s thoughts’. This could apply to forms of ecological restoration, particularly species (re)introductions where scientific predictions are used to reduce the perceived risk of unintended consequences (49). It may also apply to proposals for constructing entire systems, based on predictions that the interactions between different organisms through ‘trophic cascades’ will deliver particular outcomes (35), and is the type of control implied by Von Essen and Allen (39). This form of control must contend with an inevitable degree of uncertainty in its outcomes.

There are also ways in which conservation may attempt control by stipulating not the environment’s form but the goods and services that it is required to produce. This kind of control as ‘outputs’ could be attempted through management frameworks that require particular results, for example the legal designation of a conservation site based on the protection or expansion of particular species (36). Such bureaucratic governance, while not necessarily mandating a particular approach, may lead to attempted control through its requirement for results (50). This kind of control could apply to ‘natural capital’ approaches that measure and incentivise the delivery of particular public goods and benefits (including biodiversity indicators) (51). It could include the delivery of ‘novel ecosystems’ that, despite having no historical precedent, nevertheless produce desirable ecosystem services (52).

Each of these dimensions of control relates to Stirling’s working definition in different ways. Stabilisation attempts the realisation of a prior set of ends simply by maintaining the status quo. Control as location attempts to deliver a set of intended ends in relation specifically to geographical boundaries. Prediction attempts control, and the avoidance of negative unintended consequences, by delivering particular ends at a future point in time. Control as outputs requires the realisation of ends, defined as the products of the ecosystem in question.

These apparently different forms of potential control reveal that debates in conservation do not relate to a simple spectrum of high to low control. Rather, they are characterised by multiple different, overlapping dimensions of control. Drawing on Stirling, it is possible to attempt a working definition of control in conservation, as:

> “The realisation of a set of desired ends, and avoidance of undesired outcomes, regarding the form, location, processes and/or outputs of an ecosystem, now or in the future.”

Rewilding is held up, by some, as a means to relinquish human control of nature. Taking the definition above, it becomes important to understand whether or not rewilding marks a materially different approach to control from ‘traditional’ conservation.

### Case studies: Old Lodge and Knepp Estate

To explore these issues effectively, it is important to move beyond conceptual debates and to ground discussion in robust analysis of specific, real-world examples of practice (53). To do so, this paper compares two in-depth UK ethnographic case studies: Old Lodge nature reserve, managed by the Sussex Wildlife Trust, and the rewilding project at Knepp Estate.

Ostensibly, these two sites represent opposite ends of a spectrum of control. By exploring them in detail I aim to understand, firstly, the ways in which control operates in a ‘traditional’ UK nature reserve; and secondly, whether or not Knepp’s version of rewilding represents a significantly different approach to control, and if so in what ways.

Old Lodge, in the high weald of Sussex in southern England, comprises around 73 hectares (180 acres) and is managed as a heathland site within the wider heathland landscape of the Ashdown Forest. Management activities, undertaken by a regular volunteer group and overseen by a Sussex Wildlife Trust officer, include scrub clearance, cutting, management of bracken, and the use of grazers (both cattle and ponies). The reserve is subject to a range of formal designations, including Site of Special Scientific Interest (SSSI) and Special Protection Area (SPA), and is part of the wider Ashdown Forest Area of Outstanding Natural Beauty (AONB).

The language of Old Lodge is that of ‘high control’ conservation:

> “Sussex Wildlife Trust has managed Old Lodge Nature Reserve for over 30 years… Scrub has been removed to maintain the open heathland and woodland areas have been managed to link up and recreate areas of heathland both within the reserve and with its neighbours. Bracken has been managed by spraying and scraping… Species have responded to management well…” (54)

Knepp Estate is one of the most high profile and influential rewilding projects in the UK. Owned and run by Sir Charles Burrell and Isabella Tree, it comprises more than 1400 hectares (3500 acres) and was, until 2001, an intensive arable and dairy farm. Since then, Knepp has been managed to deliver ‘regeneration and restoration’ of nature. Knepp’s approach involves free-ranging herds of cattle, ponies, pigs and deer. The Estate sells meat from these animals, and runs nature-based tourism and camping. Knepp has gained particular attention for thriving populations of vulnerable species including nightingales (*Luscinia megarhynchos*), turtle doves (*Streptopelia turtur*) and purple emperor butterflies (*Apatura iris*). Its profile has soared since publication of the book ‘Wilding’ in 2018 (55).

Knepp is influential both for its approach to ecology and its business model – attempting to deliver financial viability through a combination of agri-environmental payments, commercial property, eco-tourism and ‘wild meat’ production. The UK is poised to transition towards a system of payment for public goods following its exit from the European Union, and Knepp was explicitly mentioned in the UK Government’s 25 Year Environment Plan as an outstanding example of ‘landscape-scale restoration in recovering nature’ (56). Consequently, Knepp is being watched closely by a wide range of landowners who see its version of rewilding as a potential model for future land management, and is an important project to understand.

In contrast to Old Lodge, the language of Knepp is of dynamic processes, uncertainty and reduced control:

> “The vision of the Knepp Wildland Project is radically different from conventional nature conservation in that it is not driven by specific goals or target species. Instead, its driving principle is to establish a functioning ecosystem where nature is given as much freedom as possible.” (57)

To understand and compare them, I spent more than 40 days at these two sites across more than a year between July 2017 and October 2018, in addition to extensive further interviews and review of written sources. At Old Lodge I joined a weekly volunteer group, accompanied employees of the Sussex Wildlife Trust to other sites, and conducted key informant interviews. At Knepp, I took part in multiple ecological surveys, meetings, conferences, tours and a safari. I was granted access to Knepp staff and participants, whom I accompanied, observed and interviewed. I spent several days accompanying the stockmen responsible for the animals, to observe Knepp’s livestock management first-hand. Taken together, this provided me with a comprehensive overview of both projects.

## 2. Ethnographic method

Ethnography originated as a method for gaining deeper understanding of (generally non-Western) cultures. In recent decades it has been used to describe a range of approaches, most commonly studies that emphasise direct observation as a primary source of information (58). It is often used in ‘case-study’ research of a system that has spatial and/or temporal boundaries and its own particular physical and cultural context (58,59). It tends to emphasise the continuous presence of the researcher in the field, though has also been applied to shorter timeframes, as for example in ‘event ethnography’ (60).

The rich context and detail provided by ethnographic studies can contribute knowledge that can only be achieved through the researcher’s experiences. This also offers the opportunity to test, and potentially falsify, existing theories and beliefs (61).

For this study I used ‘participant observation’ in which I actively took part in activities (58). This approach enabled engagement with the case studies through participation in conservation activities. It was also an appropriate way to experience how ‘insiders’ of the projects perceived and acted on issues of control that were attributed to them by themselves and others. Participant observation also enabled adjustment based on emerging ideas, lending itself to extended, qualitative research grounded in practice (62).

This research aligns with what has been described as ‘reflexive science’ (59). While some case studies may utilise ‘positivist’ methodologies that attempt to identify theoretical knowledge or generalisable facts, reflexive case studies emphasise knowledge that explicitly embraces context, providing complex, detailed and specific understanding (59). This provides knowledge that is unobtainable through ‘positive' science. It also potentially increases the risk that the researcher is influenced by the operation of power within the case study. To counter this, ethnographic researchers are encouraged to be explicit about the power dynamics involved, and to be reflexive about their own position in the inquiry.

It is possible to attempt detached and objective analysis, but with the understanding that elements of subjectivity are unavoidable in any form of scientific inquiry, not just ethnography (63). This requires reflexivity to attempt to understand the researcher’s own position in relation to the subject of study, to analyse and challenge its effects. It includes accepting that ‘truth’ can never be definitive or complete and that information may be contradictory or change over time; and continually analysing the social conditions of research, particularly the relationships between actors and between the actors and researcher, to help interpret the situations being experienced (63).

In this study, I have been conscious throughout that I come to the issue of conservation and rewilding as someone who cares personally about ‘the natural’, ‘the environment’, ‘wildlife’ and ‘wilderness’. I have a deep appreciation, aesthetically and psychologically, for both ‘nature’ and ‘wildness’. Therefore, while intellectually I am sceptical of simplistic claims or assertions about the importance of ‘nature’, I nevertheless have an emotional sympathy with arguments that value ‘nature’ and ‘wildness’ for their own sake. I am a long-term member of the Sussex Wildlife Trust, as well as an instinctive supporter of the Knepp project. I am also conscious of the prominence and influence of key individuals in this study, particularly the owners of Knepp Estate.

My broad support for the projects under consideration potentially represents ‘bias’ that I tried to bear in mind in this study. Throughout this paper, I have attempted to account for my own status as someone personally invested in the objectives of the projects being studied. In particular, I have attempted to challenge the ways in which each project is framed, where necessary, as well as my own framing of the issues. I am aware that in the process of this research, I have attempted to balance a critical social science approach, with an approach that is relevant, comprehensible, and useful for conservation practitioners and policymakers. This has led me to attempt to present the research in pragmatic terms – sometimes using language that could be interpreted as acceding to dominant, ‘mainstream’ framings of nature – but also retain high critical rigour. My hope is that this has been somewhat successful, and that this research will contribute positively to the success of conservation and the long-term status of biodiversity in the UK. By being mindful of these motivations, I have attempted at the same time to produce analysis that is of independent value.

This research was granted ethics approval by the University of Sussex Ethics Committee, reference ER/BD75/2. All participants were informed of the nature of the research. Informants received an information sheet and signed a consent form, in line with approved ethics procedure.

## 3. Results of Case Studies

### 3.1 Old Lodge

#### ‘Conservation as Control’

Many aspects of practice at Old Lodge reflect its apparent status as a ‘traditional,’ ‘high control’ conservation project: managing habitats for the benefit of particular species that are considered desirable, both through conservation and restoration.

Sussex Wildlife Trust’s management plan highlights the objective to ‘continue restoring heathland that has been invaded by Bracken, Birch and Pine’ which ‘has been the main aim at Old Lodge since inception of the LNR [Local Nature Reserve]’ (54) The management plan refers to the fact that heathland is a UK Biodiversity Action Plan priority habitat, with a ‘Habitat Action Plan for Sussex’ that aims to maintain, enhance and expand heathland with approaches including grazing management and the conversion of forestry plantations. The plan also includes Species Action Plans for particular species including nightjar (*Caprimulgus europaeus*), woodlark (*Lullula arborea*), Dartford warbler (*Sylvia undata*), silver-studded blue butterfly (*Plebejus argus*) and southern wood ant (*Formica rufa*).

Reflecting this, I frequently participated in activities designed to reduce birch (*Betula sp.*) and bracken (*Pteridium sp.*). On the site of a small disused quarry we cut birch to open the bare slopes to more light, to encourage flowering plants and insects. We pulled up small seedlings of birch and pine across the reserve, leaving only a small proportion of seedlings. The intention was to prevent the overall number of trees from increasing, and at the same time diversify their age and size distribution.

From late spring, I took part in what volunteers playfully called the ‘annual war on bracken.’ Through the summer months, a significant portion of volunteer time is devoted to cutting or pulling up bracken, often reinforced by spraying with Asulam herbicide and (in the past) large scale scraping of topsoil. This activity was described as a ‘never ending battle,’ to maintain a heathland habitat based on heather.

Areas with thick growth of birch saplings or bracken were described as ‘bad’ and the two species as ‘invasive’ and ‘bullies.’ Both species were considered a significant threat to the restoration and maintenance of the reserve. The presence of pine trees was also, to some degree, considered incompatible with the heathland status of the site (though a more complex picture of the status of trees on the site is discussed below).

Active management was the norm. On several occasions I took part in laying cut logs to block drainage channels, to improve access for people and to ‘wet up’ the area for wet-loving plants. In places, this was complemented by deliberately cutting into the moss to encourage sundews (*Drosera sp*). I took part in digging out the edges of a pond, to maintain the open water by removing grass and rush, focusing particularly on the southern edge to allow more light to reach the water, and maintaining the slope at 45 degrees to enable continued access for animals, including raft spiders (*Dolomedes fimbriatus*). At other times I took part in monitoring bird boxes, installed across the reserve for small bird species, and inspecting corrugated metal or plastic sheets left out for reptiles, including adders (*Vipera berus*).

These activities, reflecting the designation of Old Lodge as a heathland nature reserve, SSSI and SPA, suggests the presence of multiple dimensions of control. Specifically, the desire to maintain heathland and prevent vegetation succession indicates attempted ‘stabilisation.’ The focus both on increasing particular species and on the restoration of heathland generally indicates control as ‘outputs,’ while the designation of certain species as ‘invasive’ suggests elements of control as location.

**Photograph 1:**
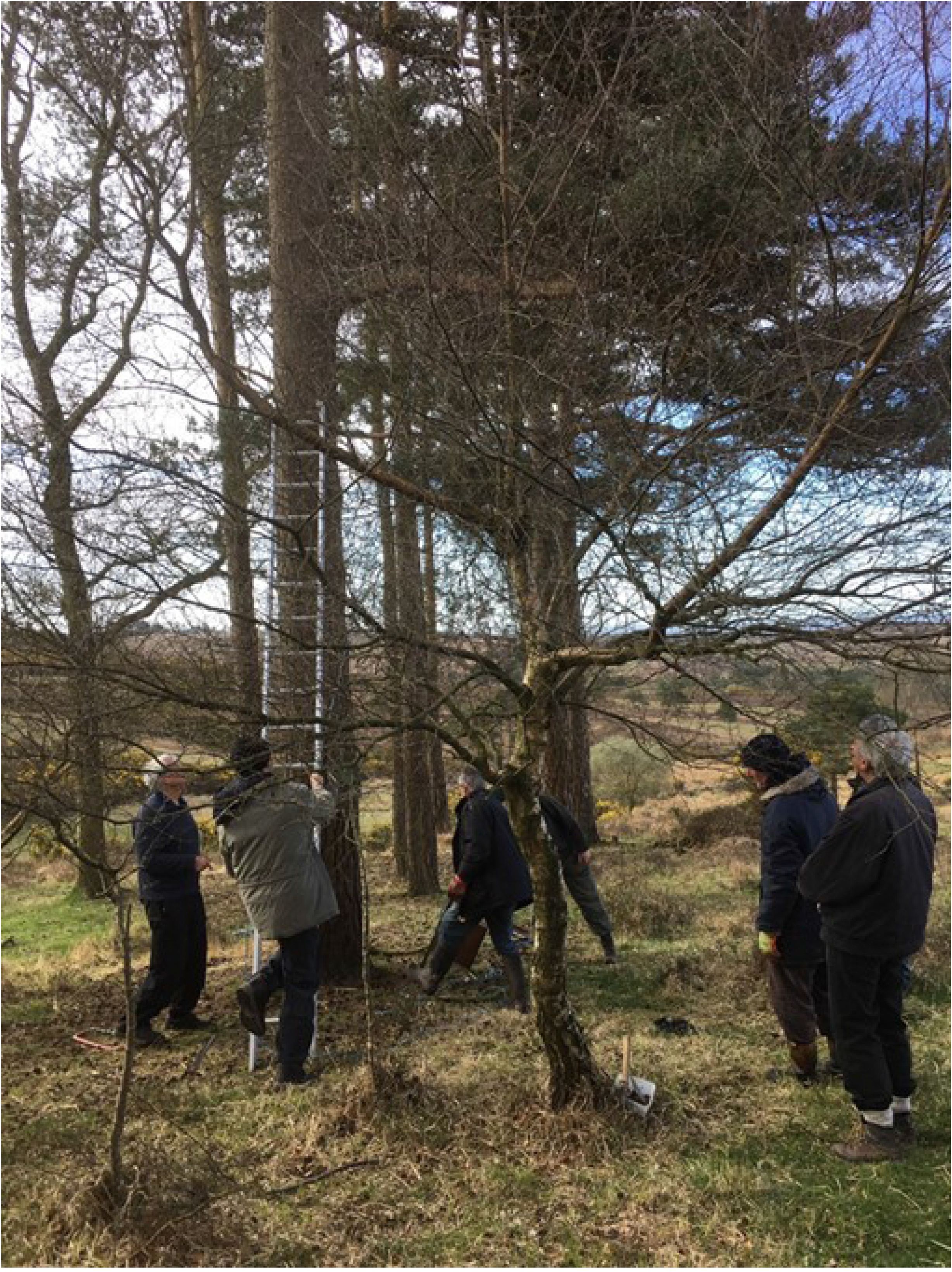
Installing a bird box at Old Lodge

#### The art of negotiated control

Within this formal approach, however, there is an element of subjectivity and flexibility. The Wildlife Trust manager responsible for the site described how the management plan is a negotiated document. It is the result of combining different perspectives, particularly from specialists in different species types, and attempting to produce a plan that delivers results in several areas.

Perhaps obviously, the nature of the plan depends on the individuals inputting into it. While the finished management plan presents activities in precise, detached and objective terms, in practice it is recognised to be much more fluid and flexible. Approaches change over time depending on changing personnel, perspectives, government frameworks and funding. The reserve manager also described how decisions about monitoring can be made to support management choices. If conservationists believe that a reserve should be managed in a particular way in future, they may choose which indicators to monitor in advance, to provide required evidence for the future management plan.

The manager draws together these strands into a plan that meets a range of different requirements, while also exercising his own judgement. He acknowledged he was not just dispassionately applying a technical approach based on ‘objective science’; there was a large degree of art involved.

Once drafted, the management plan is scrutinised by the Trust’s Conservation Committee and, as Old Lodge is an SSSI, it is submitted to Natural England for consent. This includes the question of whether it is in ‘favourable’ or ‘unfavourable’ status according to ecological indicators. While this is in principle an ‘objective’, technical issue, negotiated subjective considerations and uncertainties are apparent.

Old Lodge sits within Ashdown Forest but has a unique history that has resulted in tree cover that is considered ‘too much’ for an SSSI heathland site. Natural England therefore encourages the Trust to reduce the number of trees. Others consider the more extensive tree cover to be a valuable part of Old Lodge’s unique character and the Forestry Commission, which also has a stake in the site, would like tree cover to be maintained. This issue is therefore the subject of negotiation between multiple institutions and, for now, the site’s historical status has largely taken precedence over attempts to conform immediately with the conditions of the SSSI. The site remains in ‘unfavourable status’ as a result.

These subjective elements apparent at Old Lodge are significant when returning to the definition of control as ‘the realisation of a set of desired ends, and avoidance of undesired outcomes…’ because clearly ‘desired’ and ‘undesired’ are subjective. The use of management to maintain historical features of the site suggest elements of ‘control as stabilisation’; but this is in tension with proposed ‘control as outputs’ related to the site’s protected status. In this case, this tension is navigated not primarily with detached ecological science, but through ongoing human negotiation. Subjectivity also enters into ostensibly objective ecological indicators, with future measures of ‘control as outputs’ sometimes chosen because they are expected to provide support in advance for inherently subjective judgement calls.

#### Modern heathens

The picture of ‘control’ at Old Lodge is further complicated by the practical reality of work on the site. The volunteer leader described how much activity is guided by a sense of how the traditional commoners on the heath (the ‘heathens’) would historically have behaved: cutting trees, clearing areas and grazing animals. While these activities are consistent with the Trust’s management plan, they are not directed precisely. The volunteer group has a degree of latitude in what it does, with an ability to experiment with actions that have uncertain outcomes. Thinning trees and blocking drainage channels, for example, had a clear rationale but the longer-term consequences were unknown. While the immediate ‘outcomes’ of the work may have been precise (e.g. thinned trees) the longer term ‘ends’ were uncertain. The work was often described as ‘suck it and see’ and sometimes led to unintended results. For example, volunteers described how previous scraping of topsoil to remove bracken had resulted in bare earth being colonised by a thick growth of birch seedlings – an unwanted outcome.

Other activities, for example ‘ring-barking’ pine trees to reduce tree density but leave standing dead wood, were described as being done by ‘feel’ – a form of control but not in precise, mechanistic terms.

It is therefore not possible for the Wildlife Trust to exert total control over the site even if it wanted to, and in some cases the Trust itself attempts to reduce control. The Trust’s reserve manager described how the Old Lodge volunteers are experienced, knowledgeable and capable, but that they may be ‘too effective’ – for example being so thorough in removing tree seedlings that the site risks becoming ‘over-managed.’ Some volunteers themselves commented that their work sometimes felt like ‘gardening’ or ‘weeding’, and occasionally expressed concern that ‘there will be no trees left here at this rate.’

This presents the interesting situation in which the Wildlife Trust, potentially perceived as implementing a ‘high control’ approach to the site, is instead responsible for asking its volunteers to reduce elements of their management.

### 3.2 Knepp Estate rewilding project

Knepp’s approach is, theoretically, defined by its differences from the sort of ‘traditional’ practised at Old Lodge. As outlined above, its philosophy and language are explicitly of dynamic processes, uncertainty and reduced control. Importantly, it also represents a deliberate attempt to pioneer a new business-model for land management:

> “The aim is to show how a ‘process-led’ approach can be a highly effective, low-cost method of ecological restoration – suitable for failing or abandoned farmland – that can work to support established nature reserves and wildlife sites, helping to provide the webbing that will one day connect them together on a landscape scale.” (57)

To understand Knepp’s approach, it is important to understand both these dimensions – the philosophical and the practical – and how they combine.

Knepp’s owners are clear that the primary initial motivation for embarking on a rewilding project was financial. After inheriting the estate in 1987, they spent around 13 years attempting to make a profit through intensive arable and dairy farming, but by 1999 the business was ‘in crisis’ (Tree, 2018:31). A combination of falling market prices for milk and cereals, fluctuating agricultural subsidies, and Knepp’s low-grade clay soil, meant that despite intensification the farm made a cash surplus in only two years over a 15 year period – resulting in an overdraft of £1.5 million (Tree, 2018:39). Consequently, in 2000 the owners made the decision to abandon intensive farming and look for alternative ways to maintain the estate.

Knepp consists of three ‘blocks’ – Northern, Middle and Southern (Map 1). The management of these has differed, driven to a large extent by different sources of funding. In 2001, Knepp received Countryside Stewardship funding to restore the historic ‘Repton Park’ in the Middle block, around Knepp Castle itself, including introduction of longhorn cattle, Tamworth pigs, Exmoor ponies and fallow deer. This was extended to include the entirety of the Middle and Northern blocks in 2004, including perimeter fencing to enclose free-ranging animals. Meanwhile, reform to the EU’s Common Agricultural Policy (CAP) in 2003 enabled the Southern block to be left fallow without loss of subsidies.

**Map 1:**
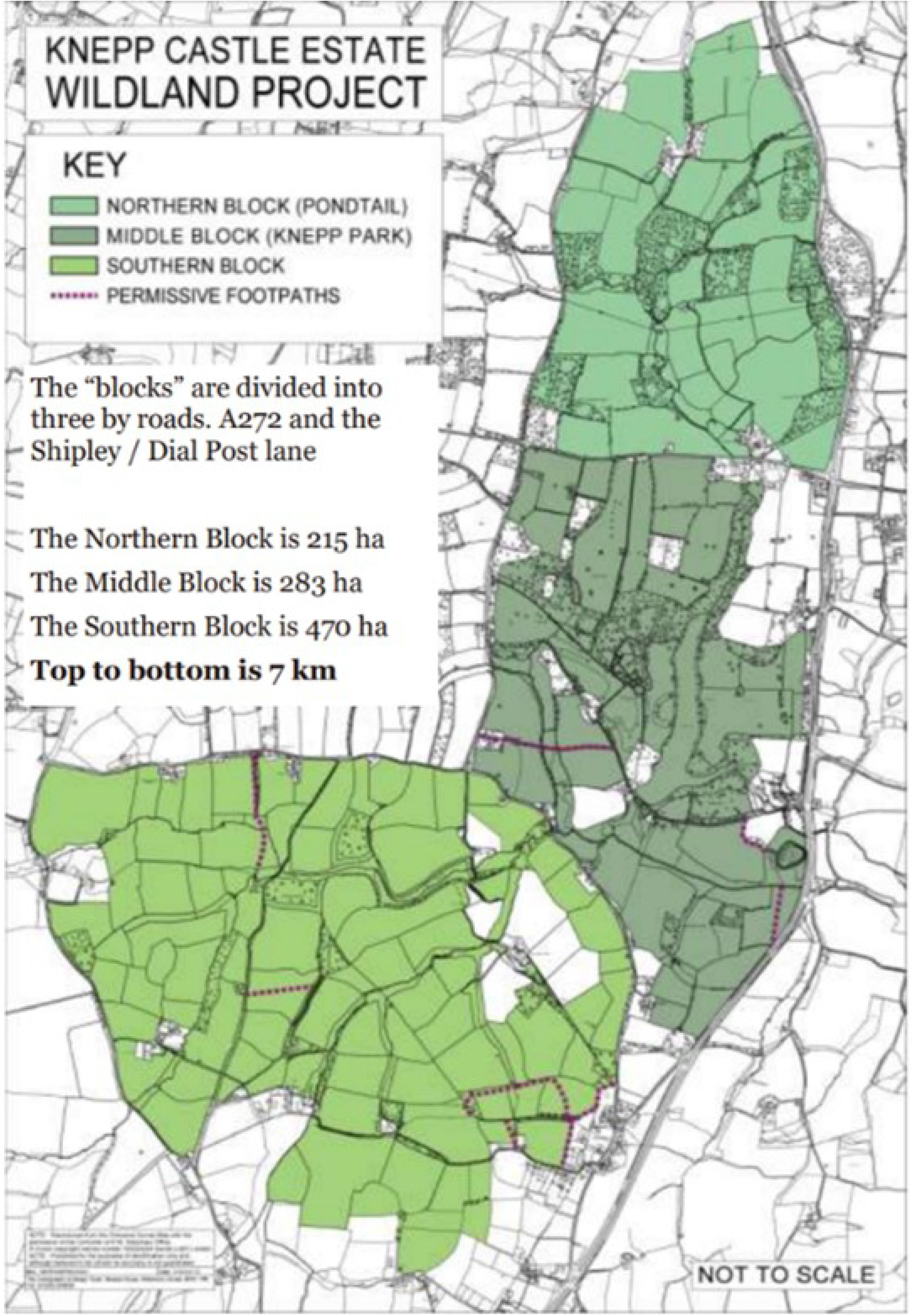
Knepp Estate (reproduced with permission from Knepp Estate)

By 2006, Knepp had developed a proposed ‘Holistic Management Plan for a naturalistic grazing project,’ but it was not until 2009 that it managed to secure Higher Level Stewardship agri-environmental funding for the whole estate. This meant the Southern block could also be enclosed to allow free-roaming animals. Unlike the Northern and Middle blocks, the Southern block therefore had six years of largely uninterrupted vegetation growth before animals were introduced – resulting in much more vigorous vegetation there now than in the other two blocks.

Knepp continues to be managed as a naturalistic grazing project, with animals – cattle, ponies, pigs and deer – enclosed within the blocks but otherwise largely free to roam. This approach was put together over time. While its progress reflected a developing philosophy of rewilding, it was equally driven by financial realities and the availability of funding.

Today, Knepp’s business model is founded on income from a range of different sources. It continues to receive agri-environmental subsidies and by relinquishing intensive farming it has released a range of buildings that have been repurposed and leased as commercial and residential properties, generating significant further revenue. The ‘wild meat’ business also generates income, as does the ‘Knepp Wildland’ camping and safari business that was established in 2014. In this sense, Knepp has not been ‘free’ to be managed according to whatever conservation approach the owners choose. It is constrained by the need for financial viability.

Despite these practical considerations, however, the rewilding philosophy underpinning Knepp is fundamental. The project is presented, first and foremost, as a conservation project, and this was borne out by the conversations I had with members of Knepp staff. For example, I was informed by interviewees that the estate could make more money from meat production if it chose to slaughter cattle at ages to optimise the grading of the beef – thereby increasing its value. However, the owner has consistently chosen to prioritise landscape management over meat production. The stockman told me:

> ‘It depends whether we put on our commercial hat or conservation hat. If commercial produce was the primary goal, then we’d manage them differently.’

The project is also committed to its rewilding philosophy in conservation terms. Initially, Knepp had the freedom to adopt a different approach because it was an intensive farm with little biodiversity value – meaning that the project began from a ‘low base’ with little to lose. Now, though, Knepp is home to large populations of threatened species, especially turtle doves and purple emperors, and I was told that some conservationists have encouraged them to begin managing the habitats more actively to support these species. These calls are being resisted. During meetings I attended, the owner said:

> ‘If you allow conservationists to push you towards managing for those species, you are still in the realms of human control.’

This position was backed up by others, with the guide on a Knepp safari saying:

> ‘Management is the most hated word. If we can do the least we can, it’s best.’

Isabella Tree describes a focus on ‘self-willed ecological processes’ and ‘restoration by letting go, allowing nature to take the driving seat’ (Tree, 2018:8).

Importantly, however, Knepp is not a value-neutral operation. The project is founded on the premise that a rewilding approach, while not aiming for specific target species, will increase the abundance and diversity of wildlife by establishing a ‘functioning ecosystem.’ Knepp’s ecologist is responsible for monitoring a wide range of species and other indicators, and reporting on them. Results that show increasing diversity and abundance are described as ‘success’.

The widespread interest in Knepp is based on such success, and this is particularly relevant regarding the UK government’s plans for future Environmental Land Management schemes (ELMS) that pay for ‘public goods,’ including biodiversity. While not focusing on individual species, Knepp’s model is required to deliver biodiversity more generally.

In terms of control, therefore, Knepp’s transition from an intensive farm to a rewilding project has altered, but not removed, a form of control as outputs. It has shifted from an exclusive focus on production of food to a primary focus on production of biodiversity. While it distances itself from ‘traditional’ conservation by eschewing target species, its model is nevertheless founded on conservation outputs. This produces tension between the presentation of Knepp as ‘low control’ and its reliance on continued production of those outputs.

#### Wood pasture

While Knepp is not focused on the delivery of target species, it is influenced by an underlying vision of a particular system – namely wood pasture.

Tree describes the profound influence on Knepp of the rewilding project at Oostvaardersplassen (OVP) in the Netherlands, pioneered by the conservationist Frans Vera (55). OVP consists of 6000 hectares of land reclaimed from the sea, designated as a nature reserve after it was colonised by large numbers of greylag geese. Cattle, ponies and deer were introduced with the intention of re-establishing natural processes and creating a ‘minimal intervention’ rewilding project (12).

OVP was founded on Vera’s proposition that much of prehistoric Europe was a mosaic environment of grassland, scrub and woodland, kept open by the activity of large herbivores that dynamically interacted with vegetation as ‘ecosystem engineers’ (64). This proposal countered earlier theories that Europe’s prehistoric landscape primarily consisted of ‘climax vegetation’ of closed-canopy forest (65).

Knepp reflects what Wynne-Jones et al (18) consider to be a general trend in UK rewilding in that it is not explicitly aiming to recreate an ecosystem from a particular point in time. Knepp’s owner told me:

> ‘You cannot recreate the past. What you’re doing is learning from the past but creating something new.’

However, the broad adherence to Vera’s theories about prehistoric ecological processes and landscapes has significantly shaped Knepp’s approach.

As at OVP, the species introduced at Knepp were deliberately chosen as proxies for extinct wild animals: longhorn cattle for aurochs; Tamworth pigs for wild boar; and Exmoor ponies for tarpan; as well as both red and fallow deer. These introductions were made with the underlying assumption that their activity would lead towards an overall result of a wood-pasture environment.

This theoretical foundation has influenced not only the initial design of the project but also its ongoing management. In particular, it manifests in the desire for a ‘balance’ between herbivores and vegetation. The owner described to me the desire to ‘get the stock numbers right in the sense of healthy, well animals and how it feels in the landscape.’ They are aiming, he said, for ‘a proper battle between animals and plants.’

This translates into practical decisions, particularly about stocking densities, based on a combination of ecological considerations and the amount of food available to the animals. The owner said:

> ‘A couple of years ago we reduced the number of animals as we felt there were too many. It is largely subjective, about a feel for the carrying capacity of the land. So we tweak the animal numbers and see what happens.’

The number of Tamworth pigs, in particular, has been reduced over time because they were found to be causing ‘too much’ disturbance and would ‘destroy everything.’ Similarly, Knepp’s owner and his stockman are required constantly to make judgements about the stocking density of cows, based on the carrying capacity of the land and their effect on the vegetation.

**Photograph 2:**
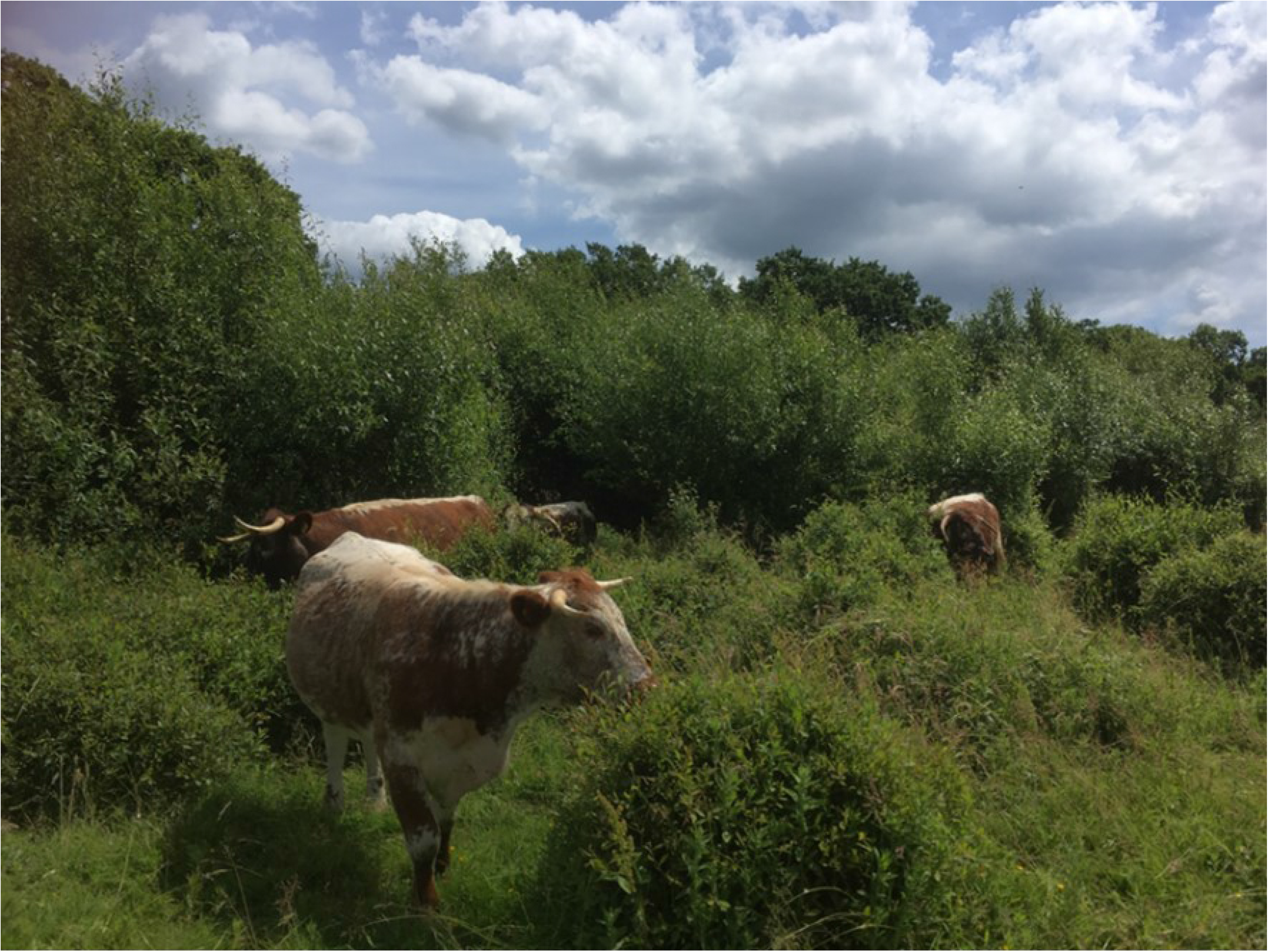
Longhorn cattle at Knepp

Here, a conscious use of language is interesting. For example, the owner considers the term ‘overgrazing’ to be too loaded, implying there is some correct level of vegetation. Other conservationists I spoke to, not employed by Knepp, have used the term in the context of managing the land to maintain a healthy, biodiverse system. It is notable that, despite seeking to establish, in the owner’s words, a ‘balanced fight’ between animals and vegetation by adjusting stocking levels, the use of the term ‘overgrazing’ is considered at Knepp to be too closely associated with farming.

Despite this, the vision of wood pasture qualifies Knepp’s overall approach of reduced control. In founding the project on the premise of a wood pasture environment, Knepp introduces elements of control through prediction (the expected results of herbivore introductions); control as outputs (the desirability of a semi-open, dynamic mosaic environment); and some degree of stabilisation (through the need for ‘balance’ in the system).

#### Animal management

Knepp’s ‘naturalistic grazing’ model, incorporating the introduction of domestic animals as proxies for extinct herbivores, means in some respects it represents a fusion of farming and rewilding – one reason it generates interest from farmers looking for a viable future business model.

A significant feature of this model is the emphasis placed on animal welfare. Here, Knepp makes a clear departure from the policy pursued at OVP, where starvation of animals has led to significant public controversy (66). In contrast, Knepp’s owner stated:

> ‘There is no appetite to have animals dying like that; it’s just not something that would work here.’

Consequently, a range of management decisions are made to ensure animals remain in good health. As outlined above, the desire to maintain a ‘healthy balance’ between animals and vegetation forms part of Knepp’s day-to-day management. This is driven in part by the underlying idea of a wood pasture environment, but also by wanting to ensure there is sufficient food for the animals.

On one occasion I took part in a fodder survey with Knepp’s ecologist. This involved assessing the availability of edible plants by estimating the extent of inedible species including fleabane, ragwort and thorny scrub. Though also of ecological interest, this exercise was primarily conducted for animal welfare reasons, including the requirement to report on the availability of fodder to the Soil Association, to maintain the organic status of Knepp’s produce. The fodder survey findings contributed to decisions on stocking levels, made by the owner and stockman based on their assessment of how much food was available.

Beyond the availability of food, ensuring high animal welfare forms an important part of the management of Knepp. On one particular day, I accompanied the stockman as he checked a calf that had not been feeding; another calf with a high temperature; a cow that had suffered an injury while giving birth; a calf with an infected ear; and a cow with an injured hip.

Over time, the policy to ensure high animal welfare has altered how Knepp is managed. When the rewilding project began the intention was for cattle, ponies and pigs to range freely throughout the year, and to reproduce in an unmanaged way. However, this presented problems. For example, cows that had trouble giving birth might go unobserved for a period of days with poor outcomes for both cow and calf. Additionally, year-old heifers would be ‘covered’ (impregnated) too young, with significant potential health impact. The owner told me:

> ‘In the wild, the heifer would die and you wouldn’t notice, but that’s not okay in this context.’

Consequently, over time more extensive management has been implemented. Now, bulls, boars and stallions are only allowed into the herds at particular times of year to ensure animals are born within a specific time window. For the cattle, this enables Knepp to bring the herds into enclosures during calving season, to monitor the health of both cows and calves and intervene if necessary – I observed a newborn calf being bottle-fed as it was not managing to suckle successfully.

The control of breeding also prevents an overpopulation of pigs that would be considered too destructive to vegetation. Additionally, with Knepp criss-crossed with footpaths and bridleways, the removal of adult males avoids potential public safety concerns presented by bulls, stallions and boars.

These decisions inevitably increase the extent of human control of the processes at Knepp – a form of control based on outputs related to animal welfare. The fact that it is a high-profile rewilding site perhaps contributes to the level of this control. Those at Knepp are anxious to avoid the animal welfare criticisms levelled at OVP, as well as any incidents involving people using the public rights of way.

Perhaps obviously, forms of human control also exist in the type of animals themselves. Though chosen as close proxies for their prehistoric wild counterparts, the cattle, ponies and pigs at Knepp are domestic species. One rewilding proponent I interviewed, not employed at Knepp, criticised the approach as perpetuating ‘farming’. In response, Knepp’s owners state that they recognise the existing limitations of the project and do not pretend that it is an unmanaged wilderness. To introduce wild boar rather than Tamworths, for example, would require a license that would not be permitted owing to Knepp’s public rights of way – an example of Knepp being constrained by external ‘control as location’ that prevents wild boar reintroduction.

**Photograph 3:**
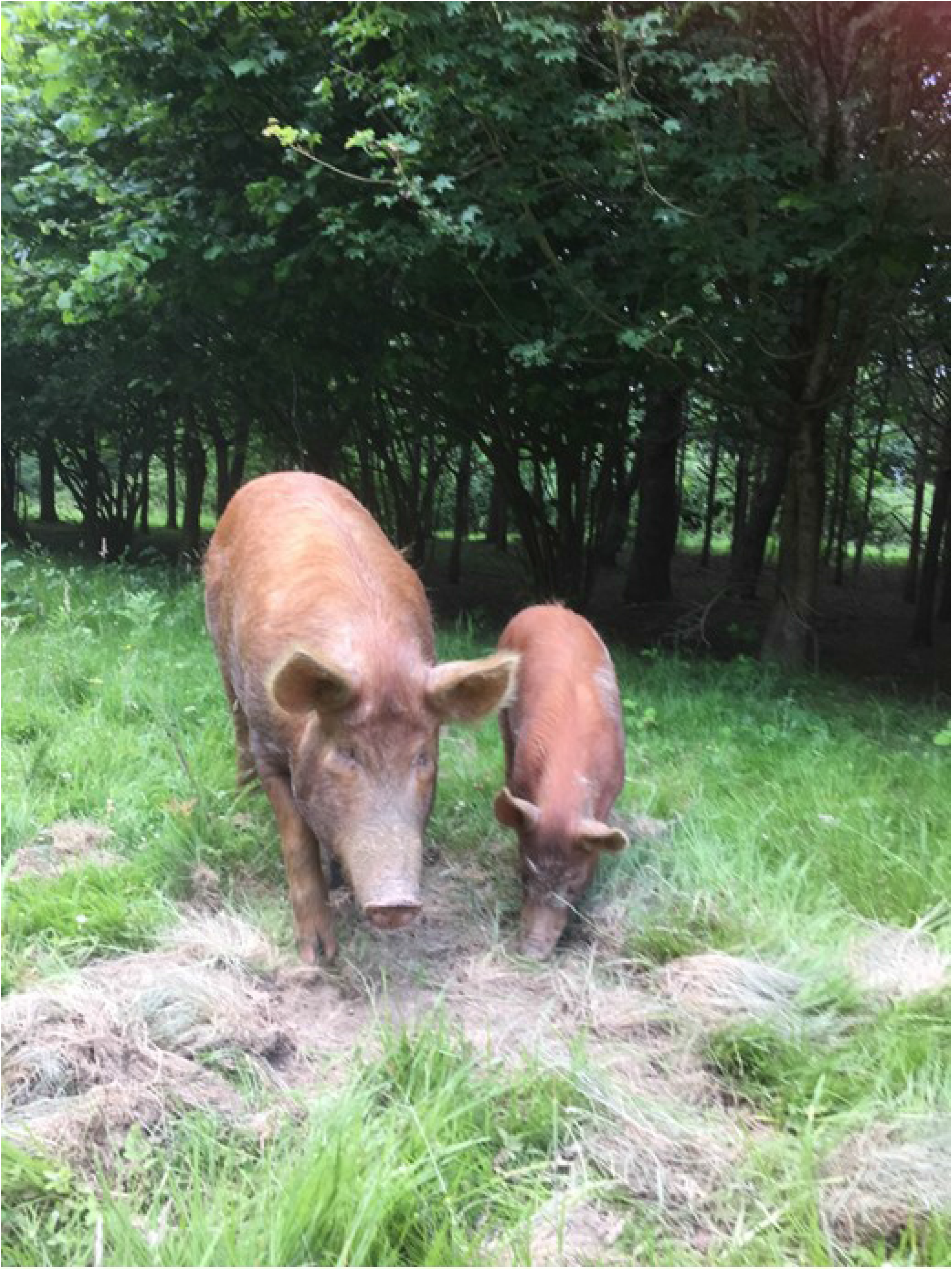
Tamworth pigs at Knepp

In many respects, therefore, while Knepp’s naturalistic grazing model is recognised as a form of rewilding, it retains aspects of farming such as animal welfare and organic certification. This limits the extent to which it can pursue ‘pure’ rewilding objectives. Nevertheless, the stockman told me that while they recognise there is a need to manage the animals, they attempt to push themselves towards lower management:

> ‘There’s a spectrum from wild to intensively farmed. We’re always trying to push towards the wild end of the spectrum, but we know it’s not possible to go the whole way.’

It is apparent that the production of ‘wild’ meat, while generating significant revenue for the Estate, is a secondary consideration to maintaining a ‘healthy’ balance between animals and the land. This suggests that, while ‘output’ forms of control are apparent, they are focused on conservation and animal welfare outputs rather than traditional farming outputs.

#### (Re)introductions

Alongside the introduction of domestic animals as proxies for extinct wild herbivores, Knepp is also undertaking introductions of wild species – specifically white storks and beavers. This important dimension of the project occupies a space within ongoing debates about (re)introductions in conservation more broadly, the implications of human interventions, the risk and uncertainty they represent, and their potential for unintended consequences (20).

The introduction of white storks at Knepp took place as part of a broader project, which aims to establish 50 breeding pairs in southern England by 2030 (67). This initially involved introducing juvenile birds from Poland into a fenced area at Knepp in 2016, with the intention that they would be ‘hefted’ to that location and return there to nest and breed as adults. In May 2020, the first white stork chicks at Knepp successfully hatched.

The introduction of white storks is presented as the restoration of an extinct native species, and therefore part of re-establishing ecological complexity and dynamism to an ecosystem that has lost many of its constituent parts. However, this project has received criticism, much of it stemming from disagreement about whether the white stork was in fact native in the recent past. Some argue that historical reports may relate to ‘vagrants,’ without storks existing in significant numbers in historical times (68). Doubts about the ‘nativeness’ of the species contribute to concerns about the effect that its release may have on existing species, including protected ground nesting birds and reptiles, with some questioning whether a sufficient Environmental Impact Assessment was carried out (69). The question of ‘nativeness’ also leads to discussion of whether such introductions represent increasing ‘wildness’ or whether in fact, if species are only present through human intervention, they represent an increased level of human management (68).

Proponents of the project argue that the white storks are historically native to southern England, and that they occupy an ecological niche that has been empty since their local extinction. They point to sightings of storks and other evidence that the species was historically present in the area, including the name of the nearby town of Storrington, meaning ‘homestead with storks’ (70). They also argue that the establishment of white storks can become emblematic of nature restoration, provide tourism opportunities and more generally reconnect people with nature (67).

This situation is relevant in a number of ways for discussion of control. In one respect, Knepp is receiving criticism for a perceived lack of control, by introducing a species that may have unknown, unintended consequences on existing wildlife. This critique would suggest an absence of control as prediction. Alongside this, Knepp invokes elements of control as location to defend the project. This includes arguing that the species ‘belongs’ in this location because it is native. The implication is that a non-native species would not be justifiable in the same way. Therefore, despite a future-facing attitude, the issue of historical nativeness is still present and active. Finally, there is a strong element of control as ‘outputs’ here: the establishment of the white stork is considered a desirable end in itself; it is also supported because of its potential to become emblematic of nature restoration, to inspire people, and to contribute overall to Knepp’s ‘success’ in increasing biodiversity.

The introduction of beavers at Knepp is different in many respects from that of storks. During my fieldwork, the question of introducing beavers was a live discussion, and since then Knepp has received approval to go ahead with an introduction, due to take place in late 2020.

In 2018 I attended a meeting at Knepp to discuss the possibility of beaver introduction. At the meeting it was clear that this could be considered controversial, and that their potential impacts would have to be mitigated to reassure local stakeholders. The meeting involved extensive discussion of the scientific research on beaver reintroduction, particularly based on examples in Devon. An expert presented the latest understanding of how beavers would behave in different conditions and what outcomes might be expected – for example, the expectation that beavers will only build dams if they do not already have access to water more than 60cm deep. The discussion also included the possibility of fencing beavers in, at least initially, to observe their behaviour before potentially letting them expand their territory. It was striking that a significant part of the discussion related to how effectively conservationists would be able to predict and/or control the beavers, should they be introduced.

Beaver introduction at Knepp therefore represents a challenge to control as stabilisation, by introducing new elements of dynamism. But in response, another form of control – ‘prediction’ – is expanded. In fact, it is precisely because Knepp is proposing to change the status quo that more extensive and precise predictions are considered necessary. Some actors questioned how realistic it is to predict the outcomes of a complex event like a species introduction; but nevertheless accepted the need to maximise the predictability of the proposals. This indicates that there is a perception that ‘society’ needs the reassurance of an element of control, despite awareness that such control can be uncertain. This chimes with analysis of beaver (re)introductions in Scotland, where the extent of beaver ‘autonomy’ has been found to be contextual and contingent (41).

Comparison of the two examples of introductions at Knepp, white storks and beavers, is instructive. The beaver project appears to deploy control as prediction to offset the loss of stabilisation. In contrast, the white stork project has been perceived as involving greater risk and uncertainty – i.e. a lack of control as prediction – and has received criticism as a result.

Similarly, a difference in control as location is visible. Both (re)introductions deploy ideas of nativeness. However, the ‘native’ status of beavers is significantly less contested than that of white storks, which has seemingly led to a greater level of acceptance for the beaver project – suggesting that forms of control as location are significant here.

## 4. Discussion

The comparison of these two case studies grounds debate about rewilding and conservation in detailed examples that add to the analysis of modern rewilding. Specifically, a focus on ‘control’ enables an in-depth exploration of whether and how an example of rewilding differs in practice from an example of ‘mainstream’ conservation.

To discuss this, it is useful to return to the working definition of control in conservation:

> “The realisation of a set of desired ends, and avoidance of undesired outcomes, regarding the form, location, processes and/or products of an ecosystem, now or in the future.”

Within this definition, control as stabilisation, location, prediction or outputs may be attempted. Table 1 outlines a broad, simplified comparison of the control dimensions between the two sites. Figure 1 visualises this same simplified analysis, illustrating different configurations of control.

**Table 1:**
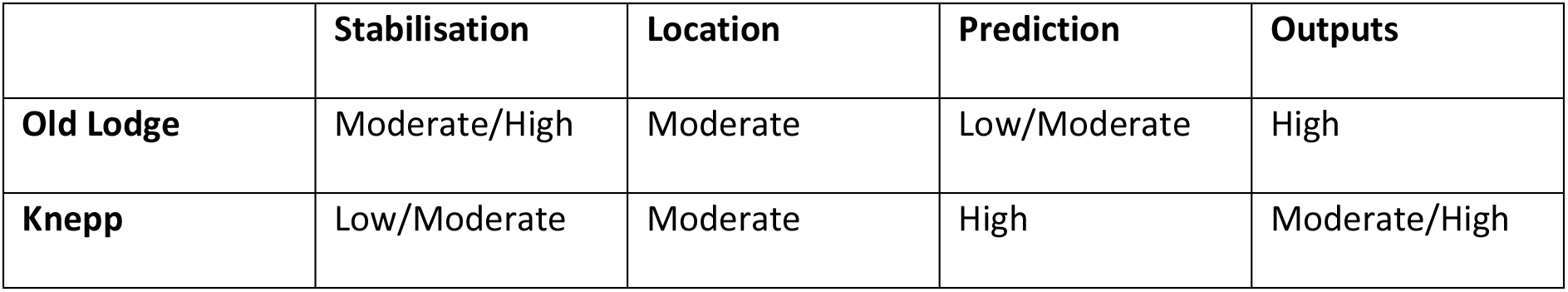
Comparison of Control Dimensions

|  | **Stabilisation** | **Location** | **Prediction** | **Outputs** |
| --- | --- | --- | --- | --- |
| **Old Lodge** | Moderate/High | Moderate | Low/Moderate | High |
| **Knepp** | Low/Moderate | Moderate | High | Moderate/High |

**Figure 1:**
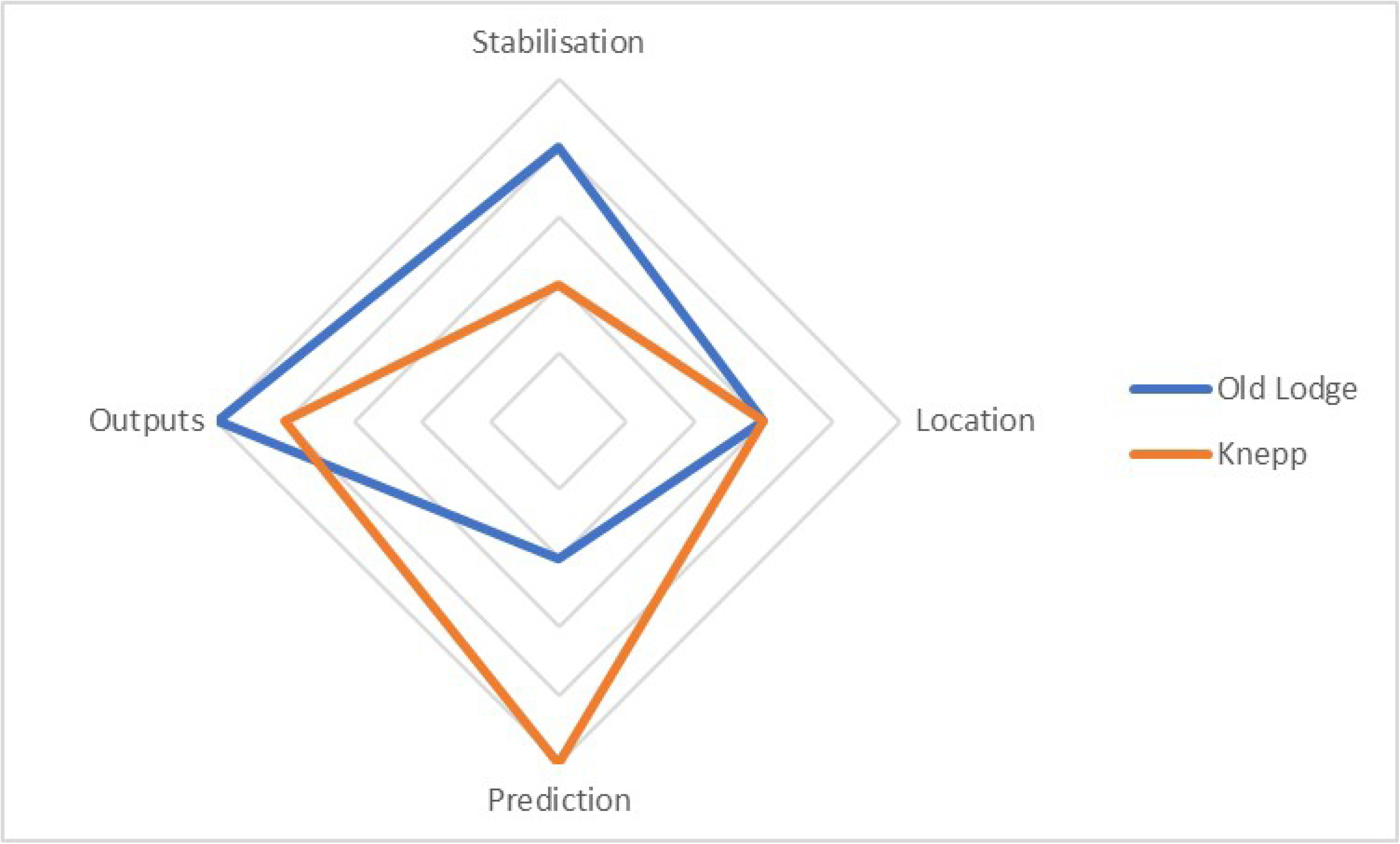
Visualisation of Indicative Configurations of Control

A range of different dimensions of control are attempted at Old Lodge. Stabilisation takes place by maintaining heathland and preventing vegetation succession, as well as the preservation of some of the historical characteristics of the site, such as tree cover. Control as outputs is apparent, both with the overall restoration of heathland and the management for particular species. Prediction is apparent in some respects – however, the existence of consistent management makes this less significant and predictions of results are often quite informal. Control as location also exists, in the sense of rejecting certain ‘invasive’ species.

Significantly, these dimensions of control are sometimes in tension with each other – for example between the stabilisation of the historical site, and the output of increasing heathland species – resulting in negotiated decisions. The formal management plan provides a foundation from which a degree of flexibility can exist. It forms parameters within which it is possible for management of the site to be done ‘by feel’ as much as by calculation. The application of control is therefore not precise. The establishment of what constitutes ‘desired ends’ is subject to negotiation and can change over time, and undesired outcomes (such as growth of birch woodland) have not always been avoided.

This underlines Stirling’s (30) observation that, for complex open-ended systems, full machine-like control is unlikely to be achieved. This understanding, in relation to a ‘traditional’ site like Old Lodge, is important for discussion of whether rewilding represents a significantly different approach.

Knepp represents an approach that has rejected defining conservation outputs as the delivery of identified species. This has not, however, removed ‘control as outputs,’ but rather shifted it. While avoiding species-based targets, the ‘success’ of Knepp’s model is reliant on a general gain in biodiversity. In addition, while the philosophy of rewilding is fundamental to the project generally, it is also integrated into the need to operate a functioning business model. The owners and their staff acknowledge that Knepp is not an example of ‘wilderness.’ Rather, it is a model for how to manage low-grade farmland in a financially viable way by producing a number of outputs, all of which are underpinned by the presentation of increased biodiversity.

Forms of control as ‘outputs’ also persist in relation to animal welfare with Knepp’s naturalistic grazing model representing, in many ways, a fusion of farming and rewilding. This is particularly evident in constant decisions around stocking densities – ensuring that there is sufficient food for the animals and avoiding both overpopulation and other health problems through control of breeding.

The management of stocking densities also represents an attempt to achieve ‘a balanced fight’ between animals and vegetation. Here, Knepp exhibits a degree of control as ‘stabilisation,’ by intervening through the adjustment of stocking densities to maintain a ‘balance’ within desired parameters.

In other respects, control as ‘stabilisation’ has been rejected, and the landscape of Knepp has clearly changed significantly from the intensive farm of 20 years ago. In place of stabilisation, however, there is evidence of increased control as ‘prediction.’ The overall design of the Knepp system is founded on the prediction that it will produce a wood pasture environment. More specifically, the planned introduction of beavers illustrates how a high level of prediction has been used to compensate for the removal of stabilisation. In contrast, the introduction of white storks has received a greater degree of criticism because of a perceived absence of prediction.

There are also elements of control as ‘location.’ To some degree this is forced upon the project: despite longer-term ambitions to expand rewilding across the landscape, currently the perimeters of the three blocks are fenced to keep the animals on-site. In other respects, control as location is apparent in the use of ‘nativeness’ to justify the introduction of both beavers and white storks, suggesting that certain species belong in the landscape while others would not.

Taken together, the observations at Old Lodge and Knepp are helpful for understanding the position of rewilding in broader conservation discourse. Rewilding is often portrayed as representing a shift from ‘high control’ to ‘low control’ – a fundamental challenge to ‘traditional’ forms of conservation. For proponents, it represents a break from ‘conservation as control’ that limits natural dynamism and impedes responses to the global biodiversity crisis. For critics, the perceived loss of control risks unintended consequences that might contribute to biodiversity loss.

The analysis presented here indicates that this characterisation is over-simplified, and that control is not a simple, linear concept. Rather, there is interplay between different dimensions of control that may become active in different ways over time and space as locations themselves, and those acting upon them, change – resulting in different configurations of control.

Old Lodge, a ‘traditional’ conservation site, negotiates a configuration of control that balances ‘stabilisation’ with ‘outputs’ – and in which control is by no means absolute. Knepp, a highly influential model of rewilding, displays reduced control as ‘stabilisation’ but potentially increased control as ‘prediction’. In comparison with Old Lodge, it represents a significant conceptual change, but not a complete departure from control. Rather, it represents a shift in the configuration of control.

These findings may reinforce the criticisms of those who argue the withdrawal of human control from ‘natural’ systems is being constrained, and that Knepp is not going far enough to relinquish human management of nature. Alternatively, this can be seen positively by those who wish to counter the perception that rewilding involves an unacceptably risky reduction in control. Instead, Knepp’s model of rewilding could be viewed as enabling a significant conceptual change without the need to abdicate control completely. This understanding has the potential to deliver greater acceptance of rewilding projects of this type.

To conclude, this analysis indicates that Knepp’s model of rewilding does represent a significant shift from the ‘traditional’ conservation of Old Lodge, opening up new conceptual spaces for conservation consistent with Wynne-Jones et al’s findings (18). By comparing the two sites, however, it becomes clear that contrary to much of the ongoing debate, discussion of rewilding should not be about whether to accept or reject ‘control’, but rather what ‘configuration of control’ is desired.

## Acknowledgements

The author wishes to thank staff and volunteers at Old Lodge Nature Reserve and the Sussex Wildlife Trust; all at Knepp Estate; and Prof. Andy Stirling and Dr Saurabh Arora for supervision and guidance. This paper does not formally represent the views of Sussex Wildlife Trust or Knepp Estate.

## References

1. Sánchez-Bayo F, Wyckhuys KAG. Worldwide decline of the entomofauna: A review of its drivers. Biol Conserv. 2019;

2. Ngo HT, Guèze M, Agard Trinidad J, Arneth A, Balvanera P, Brauman K, et al. Summary for policymakers of the global assessment report on biodiversity and ecosystem services of the Intergovernmental Science-Policy Platform on Biodiversity and Ecosystem Services [Internet]. 2019. Available from: https://ipbes.net/sites/default/files/downloads/spm_unedited_advance_for_posting_htn.pdf

3. Malhi Y. The Concept of the Anthropocene. Annu Rev Environ Resour [Internet]. 2017;42(25):1–25. Available from: https://doi.org/10.1146/annurev-environ-102016-060854

4. Crutzen PJ, Stoermer EF. The Anthropocene. IGBP Newsl. 2000;41:17–8.

5. Hamilton C, Bonneuil C, Gemenne F, editors. The Anthropocene and the Global Environmental Crisis. ABingdon: Routledge; 2015.

6. Thomas CD. Inheritors of the Earth: How Nature is Thriving in an Age of Extinction. Allen Lane; 2017.

7. Rands MRW, Adams WM, Bennun L, Butchart SHM, Clements A, Coomes D, et al. Biodiversity Conservation: Challenges Beyond 2010. 2010; Available from: http://www.sciencemag.org/cgi/content/full/329/5997/1298

8. Franks JR. Some implications of Brexit for UK agricultural environmental policy. Cent Rural Econ Discuss Pap Ser. 2016;36.

9. Jørgensen D. Rethinking rewilding. Geoforum [Internet]. 2015 Oct [cited 2016 Sep 20];65:482–8. Available from: http://www.sciencedirect.com/science/article/pii/S0016718514002504

10. Prior J, Ward KJ. Rethinking rewilding: A response to Jørgensen. Geoforum [Internet]. 2016 Feb [cited 2016 Sep 20];69:132–5. Available from: http://www.sciencedirect.com/science/article/pii/S0016718515302876

11. Sandom CJ, Dempsey B, Bullock D, Ely A, Jepson P, Jimenez-Wisler S, et al. Rewilding in the English uplands: Policy and practice. J Appl Ecol. 2019;56:266–73.

12. Lorimer J, Sandom C, Jepson P, Doughty C, Barua M, Kirby KJ. Rewilding: Science, Practice, and Politics. Annu Rev Environ Resour. 2015;40:39–62.

13. Sandom C, Donlan CJ, Svenning J-C, Hansen D. Rewilding. Key Top Conserv Biol 2 [Internet]. 2013;(February):430–51. Available from: http://doi.wiley.com/10.1002/9781118520178.ch23

14. Monbiot G. A manifesto for rewilding the world. Guard [Internet]. 2013;(May):2013–5. Available from: http://pm22100.net/01_PDF_THEMES/98_Monbiot/130527_A_Manifesto_for_Rewilding_the_World.pdf%5Cnpapers2://publication/uuid/B8E8C1E6-4423-4190-942F-251527AE3BC5

15. Rubenstein DR, Rubenstein DI. From Pleistocene to trophic rewilding: A wolf in sheep’s clothing. PNAS. 2016;113(1):E1.

16. Caro T, Sherman P. Rewilding can cause rather than solve ecological problems. Nature. 2009;462(7276):985.

17. Olwig KR. Virtual enclosure, ecosystem services, landscape’s character and the ‘rewilding’ of the commons: the ‘Lake District’ case. Landsc Res. 2016 Feb 17;41(2):253–64.

18. Wynne-Jones S, Strouts G, O’Neil C, Sandom C. Rewilding - Departures in Conservation Policy and Practice? an Evaluation of Developments in Britain. Conserv Soc. 2020 Apr 1;18(2):89–102.

19. Adams WM. When nature won’t stay still: Conservation, equilibrium and control. In: Adams W, Mulligan M, editors. Decolonizing Nature: Strategies for Conservation in a Post-colonial Era. London: Earthscan; 2003. p. 220–46.

20. Corlett RT. Restoration, Reintroduction, and Rewilding in a Changing World. Vol. 31, Trends in Ecology and Evolution. 2016.

21. Wiens JA, Hobbs RJ. Integrating conservation and restoration in a changing world. Vol. 65, BioScience. Oxford University Press; 2015. p. 302–12.

22. Hobbs RJ, Higgs E, Hall CM, Bridgewater P, Stuart Chapin III F, Ellis EC, et al. Managing the whole landscape: historical, hybrid, and novel ecosystems Manag hybrid. Source Front Ecol Environ [Internet]. 2014;12184(10223):557–64. Available from: http://www.jstor.org/stable/43187702

23. Linnell JDC, Kaczensky P, Wotschikowsky U, Lescureux N, Boitani L. Framing the relationship between people and nature in the context of European conservation. Conserv Biol. 2015 Aug 1;29(4):978–85.

24. Cronon W. The Trouble with Wilderness: Or, Getting Back to the Wrong Nature. Environ Hist Durh N C. 1996;1(1):7–28.

25. Wynne-Jones S, Clancy C, Holmes G, O’Mahony K, Ward KJ. Feral Political Ecologies?: The Biopolitics, Temporalities and Spatialities of Rewilding. Vol. 18, Conservation and Society. Wolters Kluwer Medknow Publications; 2020. p. 71–6.

26. O’Mahony K. Blurring Boundaries: Feral Rewilding, Biosecurity and Contested Wild Boar Belonging in England. Conserv Soc. 2020;18(2):114–25.

27. Marris E. Rambunctious Garden: Saving Nature in a Post-Wild World. New York: Bloomsbury; 2011.

28. Lorimer J. Wildlife in the Anthropocene: Conservation after Nature. 1st ed. Minneapolis: University of Minnesota Press; 2015.

29. Ellis E. The Planet of No Return: Human Resilience on an Artificial Earth. Breakthr J. 2012;Winter.

30. Stirling A. Engineering and Sustainability: Control and Care in Unfoldings of Modernity. In: Michelfelder DP, Doorn N, editors. Routledge Companion to Philosophy of Engineering. London: Routledge; 2019.

31. Monbiot G. Feral: Searching for enchantment on the frontiers of rewilding. Penguin Press; 2013.

32. Navarro LM, Pereira HM. Rewilding Abandoned Landscapes in Europe. Ecosystems. 2012;15(6):900–12.

33. Höchtl F, Lehringer S, Konold W. “Wilderness”: What it means when it becomes a reality - A case study from the southwestern Alps. In: Landscape and Urban Planning. 2005. p. 85–95.

34. Schnitzler A. Towards a new European wilderness: Embracing unmanaged forest growth and the decolonisation of nature. Landsc Urban Plan. 2014;126:74–80.

35. Svenning J-C, Pedersen PBM, Donlan J, Ejrnaes R, Faurby S, Galetti M, et al. Science for a wilder Anthropocene: Synthesis and future directions for trophic rewilding research. PNAS. 2015;113(4):898–906.

36. Jepson P. A rewilding agenda for Europe: Creating a network of experimental reserves. Ecography (Cop). 2016;39(2):117–24.

37. Anderson RM, Buitenwerf R, Driessen C, Genes L, Lorimer J, Svenning JC. Introducing rewilding to restoration to expand the conservation effort: a response to Hayward et al. Vol. 28, Biodiversity and Conservation. Springer Netherlands; 2019. p. 3691–3.

38. Lorimer J, Driessen C. From “nazi cows” to cosmopolitan “ecological engineers”: Specifying rewilding through a history of Heck cattle. Ann Am Assoc Geogr. 2016;106(3):631–52.

39. Von Essen E, Allen M. Wild, but Not Too-Wild Animals: Challenging Goldilocks Standards in Rewilding. 2016; Available from: http://digitalcommons.calpoly.edu/bts/

40. DeSilvey C, Bartolini N. Where horses run free? Autonomy, temporality and rewilding in the Côa Valley, Portugal. Trans Inst Br Geogr. 2019 Mar 1;44(1):94–109.

41. Ward KJ, Prior J. The Reintroduction of Beavers to Scotland: Rewilding, Biopolitics, and the Affordance of Non-human Autonomy. Conserv Soc. 2020 Apr 1;18(2):103–13.

42. Robinson DT. Control Theories in Sociology. Annu Rev Sociol [Internet]. 2007;33:157–74. Available from: www.annualreviews.org

43. Rockström J, Steffen W, Noone K, Persson Å, Chapin FS, Lambin E, et al. A safe operating space for humanity. Nature. 2009;461:472–5.

44. Marr EJ, Howley P, Burns C. Sparing or sharing? Differing approaches to managing agricultural and environmental spaces in England and Ontario. J Rural Stud. 2016;48:77–91.

45. Wilson EO. Half-Earth: Our Planet’s Fight for Life. Liveright; 2016.

46. Kopnina H. Half the earth for people (or more)? Addressing ethical questions in conservation. Biol Conserv. 2016;203:176–85.

47. Davis M. Don’t judge species on their origins. Nature. 2011;474(9):153–4.

48. Simberloff D. Non-natives: 141 scientists object. Nature. 2011;475:36.

49. Aslan CE, Aslan A, Croll D, Tershy B, Zavaleta E. Building taxon substitution guidelines on a biological control foundation. Restor Ecol. 2014;22(4):437–41.

50. Scott JC. Seeing Like a State: How Certain Schemes to Improve the Human Condition Have Failed. Yale University Press; 1998.

51. Mayer C. Valuing the invaluable: how much is the planet worth? Oxford Rev Econ Policy. 2018;35(1):109–19.

52. Perring MP, Audet P, Lamb D. Novel ecosystems in ecological restoration and rehabilitation: Innovative planning or lowering the bar? Ecol Process [Internet]. 2014 [cited 2020 Oct 6];3(8). Available from: www.ecologicalprocesses.com/content/3/1/8

53. Pettorelli N, Barlow J, Stephens PA, Durant SM, Connor B, Schulte to Bühne H, et al. Making rewilding fit for policy. J Appl Ecol. 2018 May 1;55(3):1114–25.

54. Monk-Terry M. Old Lodge Local Nature Reserve: Management Plan 2014-2024. Sussex Wildlife Trust; 2014.

55. Tree I. Wilding: The return of nature to a British farm. London: Picador; 2018.

56. UK Government. At a glance: summary of targets in our 25 year environment plan [Internet]. 2019 [cited 2019 Aug 8]. Available from: https://www.gov.uk/government/publications/25-year-environment-plan/25-year-environment-plan-our-targets-at-a-glance

57. Knepp Castle Estate. Knepp Wildland: rewilding in West Sussex [Internet]. 2020 [cited 2020 Aug 4]. Available from: https://knepp.co.uk/home

58. Gobo G, Marciniak LT. What is Ethnography? In: Silverman D, editor. Qualitative Research. 4th ed. London: Sage; 2016. p. 103–20.

59. Burawoy M. The Extended Case Method*. Sociol Theory. 1998;16(1):4–33.

60. Duffy R. What Does Collaborative Event Ethnography Tell Us About Global Environmental Governance? Glob Environ Polit. 2014;14(3).

61. Flyvbjerg B. Five Misunderstandings About Case-Study Research. Qual Inq. 2006;12(2):219–45.

62. Jorgensen DL. The Methodology of Participant Observation. In: Jorgensen DL, editor. Participant Observation. Thousand Oaks: Sage; 1989. p. 12–26.

63. Buscatto M. Practising Reflexivity in Ethnography. In: Silverman D, editor. Qualitative Research. 4th ed. London: Sage; 2016. p. 137–52.

64. Vera FWM. The Dynamic European Forest. Arboric J. 2002;26(3):179–211.

65. Ellenberg H. Vegetation Ecology of Central Europe. 4th ed. Cambridge: Cambridge University Press; 1988.

66. Kopnina H, Leadbeater S, Cryer P. Learning to Rewild: Examining the Failed Case of the Dutch “New Wilderness” Oostvaardersplassen [Internet]. International Journal of Wilderness. 2019. Available from: https://ijw.org/learning-to-rewild/

67. White Stork Project [Internet]. 2020 [cited 2020 Jun 19]. Available from: https://www.whitestorkproject.org/

68. Carter I. Bird (re)introductions: where should we draw the line? Br Birds. 2020 May;113:248–50.

69. Tout P. “Sparare Sulla Croce Rossa” - the Knepp Stork Project [Internet]. Adriawildlife. 2019 [cited 2020 Jun 19]. Available from: http://adriawildlife.blogspot.com/2019/08/sparare-sulla-croce-rossa-knepp-stork.html

70. Gow D, Campbell-Palmer R, Edgcumbe C, Mackrill T, Girling S, Meech H, et al. Feasibility Report for the Reintroduction of the White Stork (Ciconia ciconia) to England. 2016.

